# A high resolution model of the grapevine leaf morphospace predicts synthetic leaves

**DOI:** 10.1101/2024.03.08.584086

**Authors:** Daniel H. Chitwood, Efrain Torres-Lomas, Ebi S. Hadi, Wolfgang L. G. Peterson, Mirjam F. Fischer, Sydney E. Rogers, Chuan He, Michael G. F. Acierno, Shintaro Azumaya, Seth Wayne Benjamin, Devendra Prasad Chalise, Ellice E. Chess, Alex J. Engelsma, Qiuyi Fu, Jirapa Jaikham, Bridget M. Knight, Nikita S. Kodjak, Adazsofia Lengyel, Brenda L. Muñoz, Justin T. Patterson, Sundara I. Rincon, Francis L. Schumann, Yujie Shi, Charlie C. Smith, Mallory K. St. Clair, Carly S. Sweeney, Patrick Whitaker, James Wu, Luis Diaz-Garcia

## Abstract

Grapevine leaves are a model morphometric system. Sampling over ten thousand leaves using dozens of landmarks, the genetic, developmental, and environmental basis of leaf shape has been studied and a morphospace for the genus *Vitis* predicted. Yet, these representations of leaf shape fail to capture the exquisite features of leaves at high resolution.
We measure the shapes of 139 grapevine leaves using 1672 pseudo-landmarks derived from 90 homologous landmarks with Procrustean approaches. From hand traces of the vasculature and blade, we have derived a method to automatically detect landmarks and place pseudo-landmarks that results in a high-resolution representation of grapevine leaf shape. Using polynomial models, we create continuous representations of leaf development in 10 *Vitis* spp.
We visualize a high-resolution morphospace in which genetic and developmental sources of leaf shape variance are orthogonal to each other. Using classifiers, *V. vinifera, Vitis* spp., rootstock and dissected leaf varieties as well as developmental stages are accurately predicted. Theoretical eigenleaf representations sampled from across the morphospace that we call synthetic leaves can be classified using models.
By predicting a high-resolution morphospace and delimiting the boundaries of leaf shapes that can plausibly be produced within the genus *Vitis*, we can sample synthetic leaves with realistic qualities. From an ampelographic perspective, larger numbers of leaves sampled at lower resolution can be projected onto this high-resolution space; or, synthetic leaves can be used to increase the robustness and accuracy of machine learning classifiers.

**Societal Impact Statement:** Grapevine leaves are emblematic of the strong visual associations people make with plants. At a glance, leaf shape is immediately recognizable, and it is because of this reason it is used to distinguish grape varieties. In an era of computationally-enabled, machine learning-derived representations of reality, we can revisit how we view and use the shapes and forms that plants display to understand our relationship with them. Using computational approaches combined with time-honored methods, we can predict theoretical leaves that are possible to understand the genetics, development, and environmental responses of plants in new ways.

## Introduction

The field of ampelography (after the Greek *ampelos*, άμΠελος, literally vine) identifies grapevines by their phenotypic features. A focus, from the beginning, has been on leaf shape, not only because it is easily discerned, but because shape varies so dramatically between varieties. Uniquely, ampelographers quantified geometric features in leaves. When the field first arose in Europe to distinguish American rootstock varieties resistant to *Phylloxera* (Ravaz, 1902), measurements of the angles between veins, particularly those defining the petiolar sinus, were made (Goethe 1876; 1878). In the 20th century Pierre Galet used comprehensive measurements of vascular angles, dimensions, and serrations combined with extensive discussion of origin, synonyms, and other traits to catalog *Vitis vinifera* varieties (Galet, 1979; 1985; 1988; 1990; 2000). These approaches inspired further mathematical and morphometric approaches, such that mean leaves preserving details like serrations could be calculated, visualized, and used as ideal representations of varieties (Martínez and Grenan, 1999).

Our previous work has capitalized on the grapevine leaf as a morphometric model that can be used to explore the ways that genetic, developmental, and environmental influences shape complex traits (Chitwood and Sinha, 2016). Leveraging the fact that every grapevine leaf has five major veins—a midvein, two distal veins, and two proximal veins (**Fig. 1A**)—the distance between corresponding points found in every leaf can be minimized through the geometric functions of translation, rotation, scaling, and reflection. Such a superimposition of two shapes is known as a Procrustes analysis, and results from minimizing the Procrustes distance, which is a measure of overall similarity between two sets of points (Goodall, 1991). A large group of shapes can be superimposed against each other using a Generalized Procrustes Analysis (GPA; Gower, 1975). An arbitrary reference shape is selected, against which all samples are superimposed by Procrustes analysis and a new mean shape calculated. All samples are superimposed against the new mean shape and another mean is calculated. When the Procrustes distance between the mean shapes resulting from two successive iterations falls below an arbitrarily low value, the shape is taken to represent the mean against which all samples are superimposed for further analysis.

**Figure 1:**
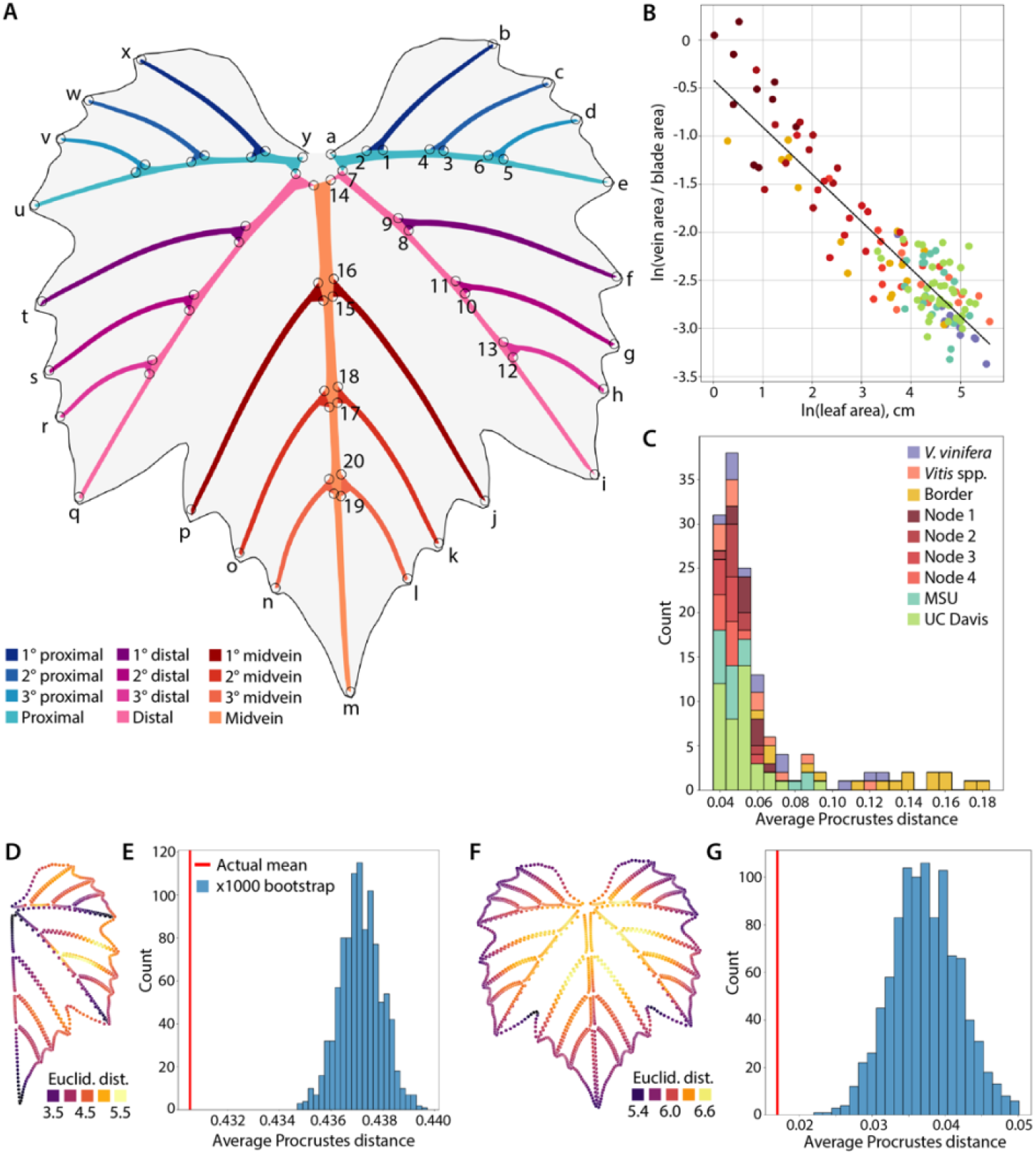
Procrustes analysis. **A)** Generalized Procrustes Analysis mean leaf. Vein identities are indicated by color. Lowercase letters indicate vein tip landmarks and numbers indicate internal vein landmarks. The numbers indicate the order the internal landmarks were detected for one side of the leaf. **B)** Negative, linear, allometric relationship between the natural log of the ratio of vein to blade area as a function of the natural log of leaf area. Data categories are indicated by color. **C)** Histogram showing the average Procrustes distance of each leaf to every other leaf. Color indicates data categories. Note that border leaves, which were purposely chosen as lying far from the center of data, have higher average Procrustes distances than other leaf types. **D)** The average Euclidean distance of each point to its counterpart between leaf halves superimposed using Procrustes analysis. **E)** The actual mean and a distribution of 1000 bootstrap simulations of the Procrustes distance of each leaf half to its counterpart. **F)** The average Eulcidean distance of each point to its counterpart between leaf pairs from 20 varieties. **G)** The actual mean and a distribution of 1000 bootstrap simulations of the Procrustes distance of each leaf to its counterpart for the 20 pairs.

Over the last decade, we have successively increased the resolution of our morphometric techniques, the number of leaves we study, and the axes of genetic, developmental, and environmental variation which we measure that modulate leaf morphology (**Table 1**). We began by examining the genetic basis of leaf shape between *Vitis vinifera* varieties in the United States Department of Agriculture (USDA) Agricultural Research Services (ARS) National Clonal Germplasm Repository at Wolfkill (Winters, California, USA), using 10 landmarks across both sides of the leaf (Chitwood et al., 2014). We subsequently analyzed leaves from the USDA Cold-Hardy Grape (*Vitis*) Collection in Geneva (New York, USA), which in contrast to Wolfskill contains mostly North American and Asian *Vitis* spp., and for which we collected leaves from an entire shoot as a developmental series (which was not done for the Wolfskill leaves) and measured using 17 landmarks across the whole leaf (Chitwood et al., 2016a). In follow up work we increased the landmark number to 21 to only half of the leaf, marking the bases of the veins that allowed us to measure the area occupied by vasculature relative to blade and to observe the exponential decrease in the ratio of vein-to-blade area as leaves expand during development (Chitwood et al., 2016b; Chitwood et al., 2021). We next created continuous developmental models of leaf shape, modeling each *x* and *y* coordinate value as a polynomial function of node position in the shoot and using the resulting “composite” leaf shapes to show that they discriminate species identity better than individual leaves (Bryson et al., 2020). Using 21 landmarks on the Geneva leaves and in comparison to *V. vinifera* leaves derived from California breeding populations, we calculated a morphospace: a continuous representation that can be used to visualize and demarcate the extent of genetic, developmental, and environmental variation in the genus *Vitis* (Chitwood and Mullins, 2022).

**Table 1:** Previous ampelographic work and leaves analyzed.

| Publication | Chitwood et al., 2014 | Chitwood et al., 2016a | Chitwood et al., 2016b | Bryson et al., 2020 | Chitwood, 2021 | Chitwood and Mullins, 2022 | This manuscript |
| --- | --- | --- | --- | --- | --- | --- | --- |
| Landmarks | 10 | 17 | 21 | 21 | 24 | 21 | 90 |
| Pseudo-landmarks | 0 | 0 | 0 | 0 | 5999 | 0 | 1672 |
| Whole or half leaf | whole | whole | half | half | half | half | whole |
| Vein-to-blade ratio | no | no | yes | yes | yes | yes | yes |
| Developmental modeling | no | no | no | yes | no | yes | yes |
| Predicted morphospace | no | no | no | no | no | yes | yes |
| No. leaves Wolfskill, CA | >9500 | 9548 | 0 | 0 | 240 | 0 | 14 |
| No. Leaves Geneva, NY | 0 | 3292 | >5,500 | 8465 | 0 | 6284 | 66 |
| No. Leaves UC Davis, CA | 0 | 0 | 0 | 0 | 0 | 0 | 59 |

The above studies rely on a small number of homologous landmark points to represent leaf shape. The intricate curves and details of leaves that captivate our eyes are missing. By placing a high number of pseudo-landmarks—equidistant points along line segments connecting landmarks—we can analyze the remaining details that comprise leaf morphology. This approach uses tracing and is more time-consuming and results in a different strategy: instead of measuring thousands of leaves at a resolution of dozens of landmarks, hundreds of leaves are measured using thousands of pseudo-landmarks. Previously we measured half the leaf of representative wine and table grape varieties (Chitwood, 2021). Here, we extend high resolution morphometric analysis to the entire leaf across both *V. vinifera* varieties as well as wild *Vitis* spp., and for the first time measure leaves from rootstock varieties in the Teaching Vineyard at the Robert Mondavi Institute for Wine and Food Science at University of California at Davis (UC Davis). We create continuous developmental models for ten *Vitis* spp. and project these models onto a visualized morphospace. Classifiers accurately predict genotype and developmental identities of leaves. Using models, we classify leaves from the morphospace and use it to create synthetic leaves: high resolution leaf representations that are theoretically possible based on the known borders of the *Vitis* morphospace. We end with a discussion about how synthetic leaves can be incorporated into workflows to improve studies of leaf shape using machine learning approaches.

## Materials and Methods

### Plant material and data collection

Leaves from *V. vinifera* varieties, rootstock varieties, and wild *Vitis* spp. were sampled from the Teaching Vineyard at the Robert Mondavi Institute for Wine and Food Science at UC Davis. Original scans with scale for each leaf sampled from the UC Davis teaching vineyard are available (Chitwood, 2024). Leaves from the USDA Wolfskill, CA and Geneva, NY collections were measured by selecting leaves from previously published data (Chitwood, 2020; Chitwood et al., 2020).

Leaves were categorized in two different ways. In the first, leaves were factored by how they were measured and the intention of including them in the data. The factor levels and their descriptions are: “*V. vinifera*”, representative leaves from *V. vinifera* varieties in the Wolfskill collection; “*Vitis* spp.”, representative leaves from wild *Vitis* spp. in the Geneva collection; “Border”, leaves from both the Wolfskill and Geneva datasets that were sampled because they lie far from the center of the morphospace; “Node 1”, “Node 2”, “Node 3”, and “Node 4” leaves which represent developmental series from ten wild *Vitis* spp. from the first four nodes counting from the growing tip; “MSU” and “UC Davis”, which are leaves traced by students at Michigan State University and University of California at Davis, respectively, from the UC Davis Teaching Vineyard. In the second factorization, leaves are classified by their genetic or developmental identities. Each leaf has exactly one assigned genetic and development factor level. Genetically, leaves are classified as arising from “*V. vinifera”*, “*Vitis* spp.”, “Rootstock”, or “Dissected” types. Developmentally, leaves are classified, as counting from the growing tip, as arising from “Node 1”, “Node 2”, “Node 3”, “Node 4”, or as “Mature” if greater than node 4.

### Automatic landmark detection

For each leaf, starting on one side of the petiolar junction (point “a” or “y” in **Fig. 1A**), in ImageJ (Schneider et al., 2012) or Fiji (Schindelin et al., 2012) a trace of the vein and blade was made using the line segment tool and stored as a .txt file. An information file, storing species or variety name, the dataset the leaf scan originated from, the vine, developmental stage, file name of the associated image, dataset source, and scaling information, was stored as a .csv file. All three files have a three digit identifier indicating if they are from the Wolfskill or Geneva datasets (starting with “0”) or the UC Davis vineyard dataset traced by MSU (starting with “1”) or UC Davis (starting with “2) students separated by underscore and followed by the species or variety name in capital letters. Another underscore separates an identifier indicating if the file is a blade or vein trace or information file (“blade”, “vein”, and “info”, respectively). File names from a data folder were read in and used to store raw data and associated metadata. All analyses were performed in Python (version 3.10.9), Numpy (Harris et al., 2020), and Pandas (McKinney and the Pandas Development Team, 2015) using a Jupyter notebook (Kluyver et al., 2016). Matplotlib (Hunter, 2007) and Seaborn (Waskom, 2021) were used for data visualization.

90 landmarks (25 from the vein tips, 40 internal vein landmarks, and 25 blade landmarks) were automatically detected. Vein and blade traces were interpolated with 1000 equidistantly spaced points to increase the resolution of the data using the interp1d function SciPy (Virtanen et al., 2020). The petiolar junction was calculated as the mean of the first (“a) and last (“y”) coordinate in the vein trace file. The Euclidean distance of each vein trace point to the petiolar junction was calculated as a function of geodesic distance along the trace. The SciPy find_peaks function was used to find the 25 vein tip indices in the trace using the above plot. The 25 blade landmarks correspond closely to the 25 vein tip landmarks, and are identified by index as the closest point in the interpolated blade trace to an identified vein tip landmark. The 25 vein tip landmarks are labeled “a” through “y” in **Fig. 1A**.

40 internal landmarks, placed on each side of the base of a vein, were calculated in the order indicated in **Fig. 1A**. The landmark on the more distal side (farthest away from the petiolar junction) of the base of a vein was calculated before the proximal side (closer to the petiolar junction). The first calculated internal landmark of the base of a vein was used in reference to calculate the other. Internal landmarks were calculated by specifying a start and end index along a segment of the vein trace and a reference index. Calculating the Euclidean distance for each point along the segment from start to end to the reference index, the index with either a maximum or minimum distance value was assigned as the landmark. For example, internal landmark “1” was calculated using landmark “b” as the start index, landmark “c” as the end index, and landmark “c” as the reference index (**Fig. 1A**). Internal landmark “1” was calculated as the index with the maximum Euclidean distance along this segment from landmark “c”.

Landmark 2 was calculated using landmark “a” as the start index, landmark “b” as the end index, and internal landmark “1” as the reference index. Internal landmark “2” was calculated as the index with the minimum Euclidean distance along this segment from landmark “1”. The rest of the internal landmarks were calculated in a similar way, specifying the segment of the trace using previously calculated landmarks on which they fall, and calculating the minimum or maximum Euclidean distance from a reference landmark.

Along each line segment of the vein trace defined by internal and tip landmarks, and each line segment of the blade trace defined by blade landmarks, 20 equidistant pseudo-landmarks were calculated. Redundant landmarks at the beginning and end of each segment were eliminated. There were 1672 pseudo-landmarks total: 1216 vein and 456 blade.

### Vein-to-blade ratio, Procrustes analysis, and biological reproducibility

The ratio of the vein area to the blade area (vein-to-blade ratio) was calculated using Gauss’ area formula (or, the shoelace algorithm; Meister, 1769; Braden, 1986). Vein-to-blade ratio is an allometric indicator with a negative, linear relationship to leaf area (Chitwood et al., 2021). The vein and blade pseudo-landmarks were treated as vertices of a polygon from which area was calculated, which were used as the area of the venation and leaf, respectively. The blade area was calculated by subtracting the vein area from the leaf area. The natural log of vein-to-blade ratio was plotted against the natural log of leaf area to which a line was fitted using the SciPy curve_fit function.

The mean Procrustes distance of each leaf to every other leaf was calculated using the procrustes function from SciPy and plotted as a histogram. To assess the ability of our morphometric method to measure biological reproducibility, we compared the two halves of each leaf to each other as well as a set of 20 pairs of leaves originating from the same variety in the UC Davis collection and calculated Procrustes distances. For each of the two analyses, 1000 bootstrap calculations of the average Procrustes distance were calculated by randomizing one set of the data against the other and comparing the distribution to the actual average. For each pseudo-landmark from each comparison, a Euclidean distance was calculated and projected back onto Generalized Procrustes Analysis mean leaf to visualize the spatial contribution of each pseudo-landmark to the alignment.

### Developmental modeling, Principal Component Analysis, and Linear Discriminant Analysis

Developmental modeling was undertaken using the same techniques as Bryson et al. (2020). For 10 wild *Vitis* spp., using the first four expanded leaves from the growing tip, for each of 3344 *x* and *y* coordinate values from 1672 pseudo-landmarks, a second degree polynomial model was fitted as a function of node number counting from the tip (1, 2, 3, 4) using the Numpy polyfit function. 100 leaves were modeled from node 1 to 4 and used in subsequent analyses.

The PCA function from Scikit-learn (Pedregosa et al., 2011) was used for Principal Component Analysis (PCA). Eigenleaf representations were calculated using the inverse_transform function. The LinearDiscriminantAnalysis function from Scikit-learn was used for Linear Discriminant Analysis (LDA). The predict function was used on resulting models to classify leaf identities. The confusion_matrix function was used to calculate and visualize confusion matrices. To calculate a convex hull in a 9 dimensional principal component space, the ConvexHull function from SciPy was used followed by the Delaunay function to find simplices. A uniform distribution of points was sampled using the Dirichlet distribution using the SciPy dirichlet function.

## Results

Using automatic landmark detection, continuous representations of the size and shape of leaves identifying specific veins can be visualized, allowing differences in leaf morphology between *V. vinifera* varieties and *Vitis* spp. (**Fig. 2A**), atypical leaves bordering the known *Vitis* morphospace (**Fig. 2B**), developmental series of leaves from the first four nodes at the growing tip of ten *Vitis* spp. (**Fig. 2C-L**), and *V. vinifera* and rootstock variety leaves from the UC Davis collection traced by MSU (**Fig. 3A**) and UC Davis (**Fig. 3B**) students to be qualitatively compared at a glance. In addition to visual inspection, we used a number of other approaches to verify the quality of our input data. As previously shown for all *Vitis* spp. leaves (Chitwood et al., 2021), our leaves follow a negative linear relationship between the natural log of the ratio of vein-to-blade area as a function of the natural log of leaf area (**Fig. 1B**). Calculating the average Procrustes distance of each leaf to all other leaves, border leaves had a higher distance from other leaves, which is expected as they were purposely chosen to lie far from the center of the morphospace (**Fig. 1C**). To determine if our method could detect biological reproducibility, we compared the Procrustes distances of 1) each half of every leaf to the other (**Fig. 1D-E**) and 2) twenty pairs of leaves from the UC Davis collection that originate from the same variety (**Fig. 1F-G**). In both cases, the actual mean Procrustes distance was less than x1000 bootstrap simulations.

**Figure 2:**
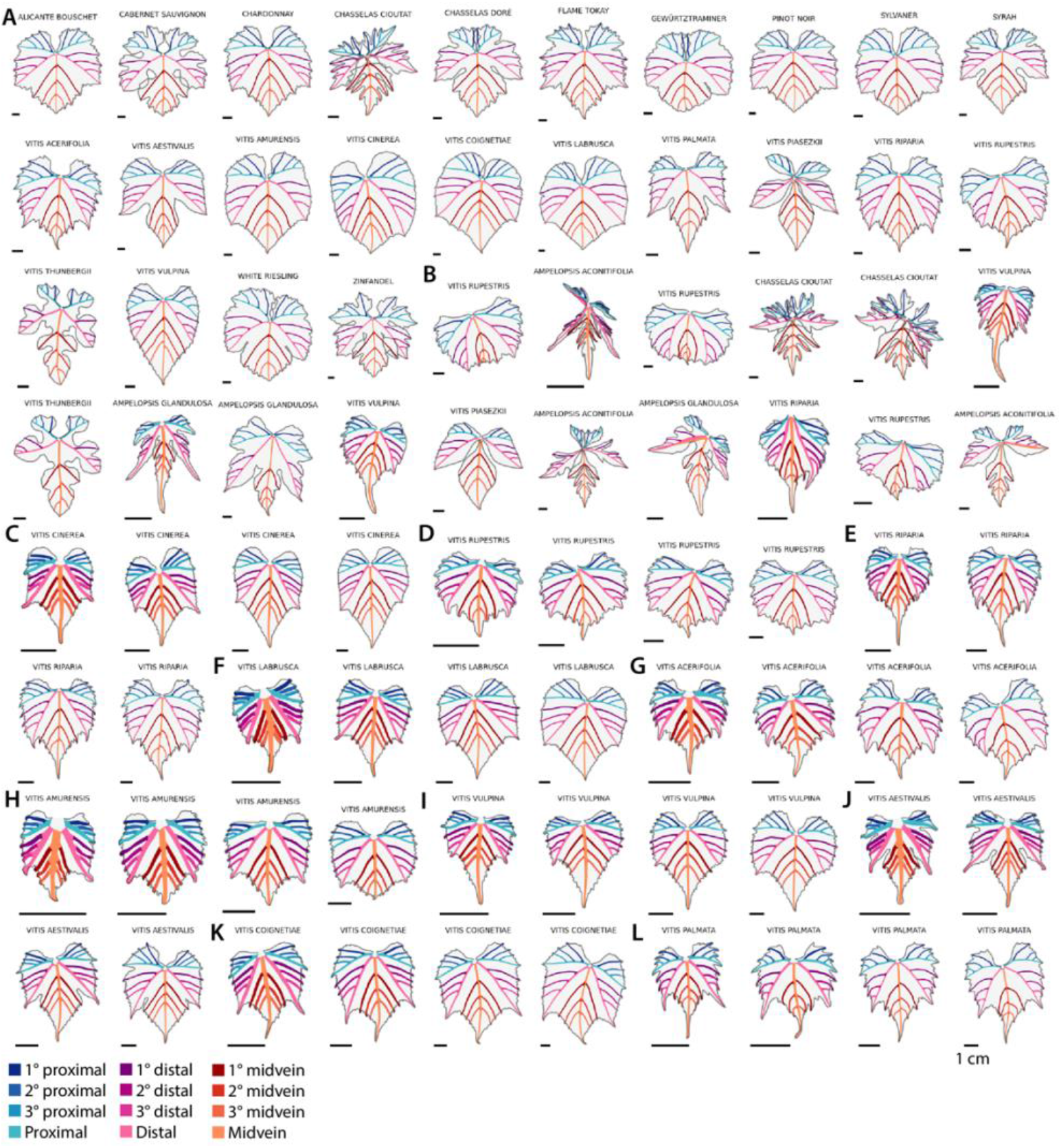
Leaf trace data. Original trace data for each leaf. Vein identities are indicated by color and 1 cm scale bar is provided. Leaves are arranged by data category, left-to-right, top-to-bottom. Species or variety name is indicated. **A)** *V. vinifera* and *Vitis* spp. leaves. **B)** Border leaves. **C-L)** Developmental series leaves. Leaves are grouped in fours counting from the growing tip: node 1, node 2, node 3, and node 4. Developmental series for 10 species were measured: C) *V. cinerea*, D) *V. rupestris*, E) *V. riparia*, F) *V. labrusca*, G) *V. acerifolia*, H) *V. amurensis*, I) *V. vulpina*, J) *V. aestivalis*, K) *V. coignetiae*, and L) *V. palmata*.

**Figure 3:**
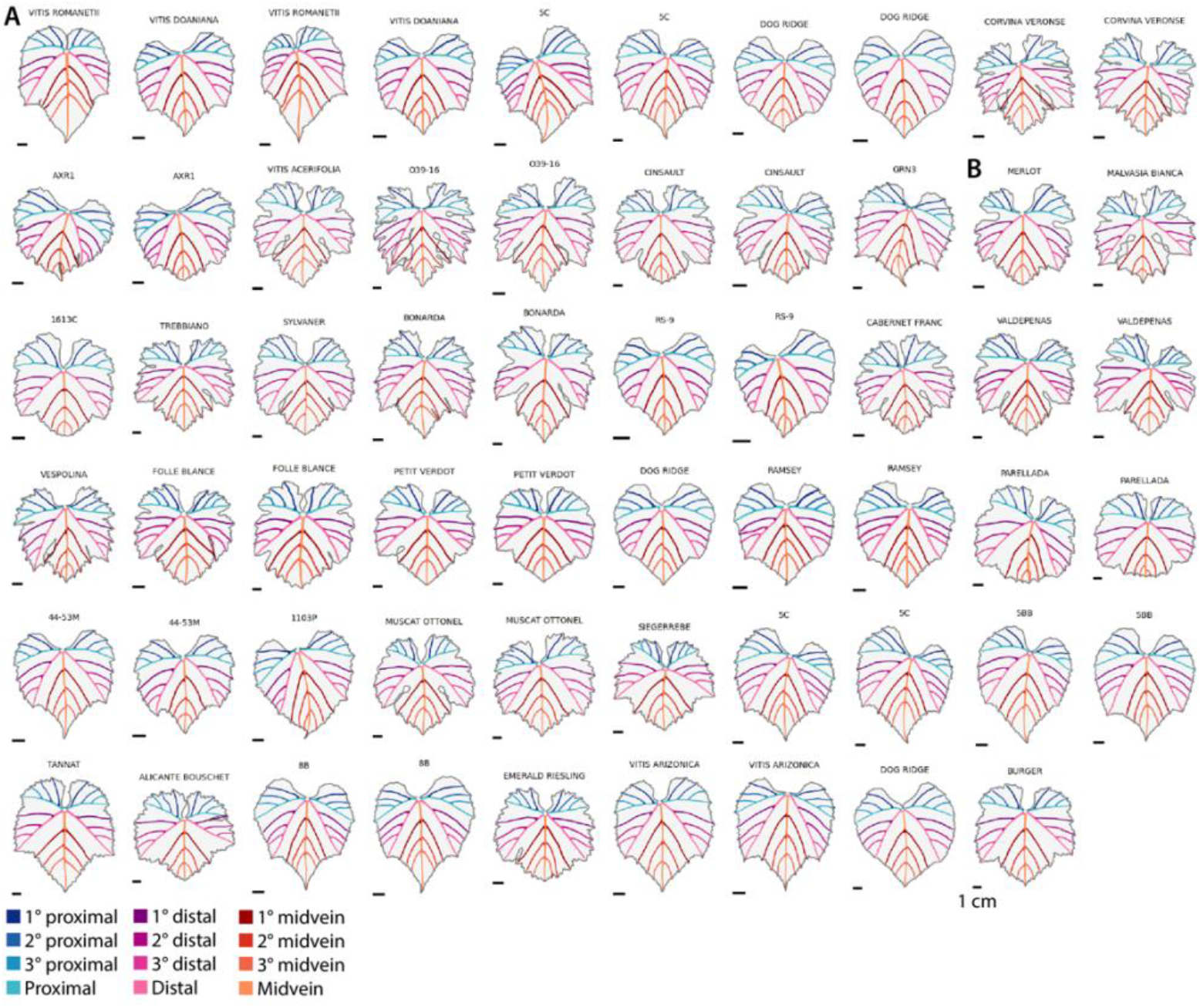
Leaf trace data. Original trace data for each leaf. Vein identities are indicated by color and 1 cm scale bar is provided. Leaves are arranged by data category, left-to-right, top-to-bottom. Species or variety name is indicated. **A)** Leaves traced by Michigan State University students and **B)** leaves traced by University of California at Davis students.

Comparing Euclidean distances of superimposed leaf halves, it was the leaf tips that more closely aligned with each other and contributed to the biological reproducibility (**Fig. 1D**), whereas comparing leaf pairs originating from the same variety, it was the overall blade outline, but especially the vein tips and the distal sinuses (**Fig. 1F**).

For ten *Vitis* spp. we modeled each *x* and *y* coordinate value across the first four nodes from the growing tip of the vine as polynomial functions, creating continuous representations of leaf development (**Fig. 4A**). Decreases in vein-to-blade ratio value are observed across each series, which allometrically reflects increases in leaf size during development. To visualize these models in the context of shape variation represented among all the measured leaves, we performed a Principal Component Analysis (PCA), which not only is a dimension reduction technique that allows more variation to be analyzed using fewer axes, but also through an inverse transformation, allows the underlying morphospace to be visualized (based on “eigenleaf” representations). The first principal component explains 46.9% of total variation and is associated with shape variation ranging from a *V. rupestris*/*V. riparia*-like leaf shape with a shallow petiolar sinus and reduced lobing to a more *V. vinifera*-like leaf shape with an overlapping petiolar sinus and deeper lobes (**Fig. 4B**). PC2, explaining 10.6% of total variation, shows a more dynamic range of vein-to-blade ratios and ranges from a developmentally mature, orbicular leaf type with no lobing and deep petiolar sinus to a developmentally younger leaf type with a shallow petiolar sinus and pronounced midvein that is relatively long. Although somewhat confounded, as previously described (Chitwood and Mullins, 2022) axes explaining genetic differences between species and varieties vs. developmental differences are orthogonal to each other. Genotype differences (between *V. vinifera, Vitis* spp., rootstock, and dissected classes) mostly vary across PC1 and diagonally from low PC1/PC2 to high PC1/PC2 values (**Fig. 4C**). Contrastingly, developmental differences vary from small, young leaves (high vein-to-blade ratio) to large, mature leaves (low vein-to-blade ratio) across PC2 and diagonally from low PC1/high PC2 to high PC1/low PC2 values. If continuous developmental models of leaves (**Fig. 4A**) are projected onto the morphospace (**Fig. 4D**), they confirm the direction of developmental trajectories.

**Figure 4:**
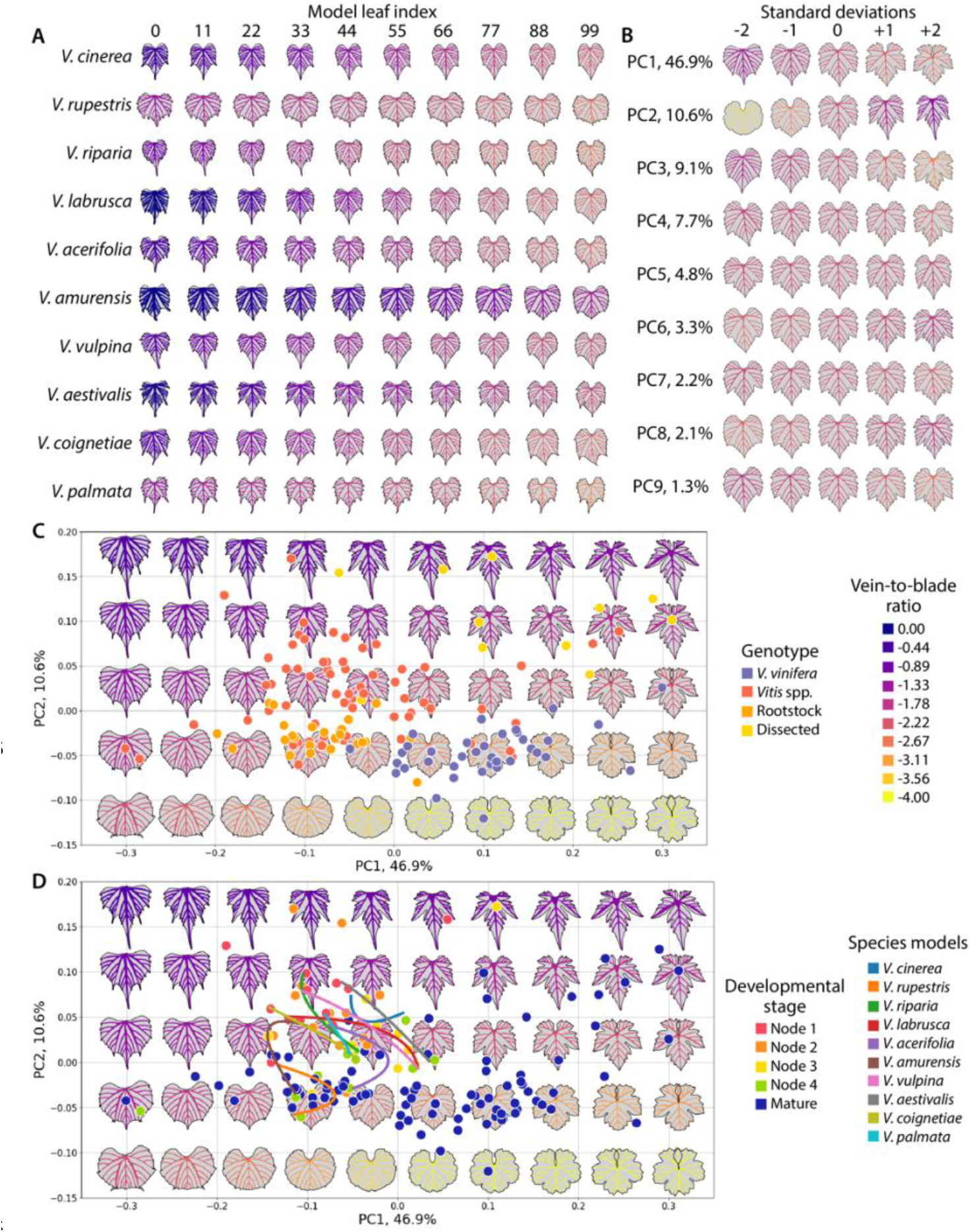
Morphospace. **A)** Continuous models of leaf development for 10 *Vitis* spp. From node 1 to node 4 counting from the growing tip, 100 leaves were modeled, for which 10 equally spaced indices are shown. **B)** Eigenleaf representations at -2, -1, 0, +1, and +2 standard deviations and percent variance explained for the first 9 principal components. **C)** Morphospace onto which data colored by genotype is projected. **D)** Morphospace onto which data colored by developmental stage is projected. Developmental models for 10 species, which are not included in the calculation of the morphospace, are projected onto it as curves, indicated by color. Values of the natural log of the ratio of vein to blade area are consistent between panels and figures, as indicated.

The orthogonality of genotypic and developmental variation suggests that these axes can be independently predicted from each other. We created classifiers for leaves based on genotypic (**Fig. 5**) and developmental (**Fig. 6**) identity using Linear Discriminant Analysis (LDA) which we then used to predict the identities of theoretical morphospace leaves. Average leaf shapes across genotype classes (remembering that *Vitis* spp. leaves contain developing leaf shapes from nodes 1-4 not represented in other classes) mostly vary by the degree of lobing (**Fig. 5A**). Except for 3 leaves, all leaves are correctly classified by their genotypic class (**Fig. 5B**) and show separation in the LDA space (**Fig. 5C-D**). If 200 theoretical eigenleaves from the morphospace are synthesized and classified, the separation of genotype classes across low PC1/PC2 to high PC1/PC2 values is evident. Contrastingly, variation across average leaf shapes by developmental node (remembering that nodes 1-4 are mostly represented by *Vitis* spp.) is more gradual, with a shallow-to-deeper petiolar sinus and a midvein which shortens in relative length as leaves mature and grow in size, in addition to strong reductions in the ratio of vein-to-blade area (**Fig. 6A**). All but one leaf is correctly predicted by its developmental class (**Fig. 6B**). Leaves from different nodes separate by class in LDA space (**Fig. 6C-D**), but especially by LD2 vs. LD1, and are continuously connected to each other when developmental models are projected onto the space (**Fig. 6C**). If eigenleaf representations are classified by developmental stage, a gradient from small and young leaves to large and mature leaves is observed along low PC1/high PC2 to high PC1/low PC2 values.

**Figure 5:**
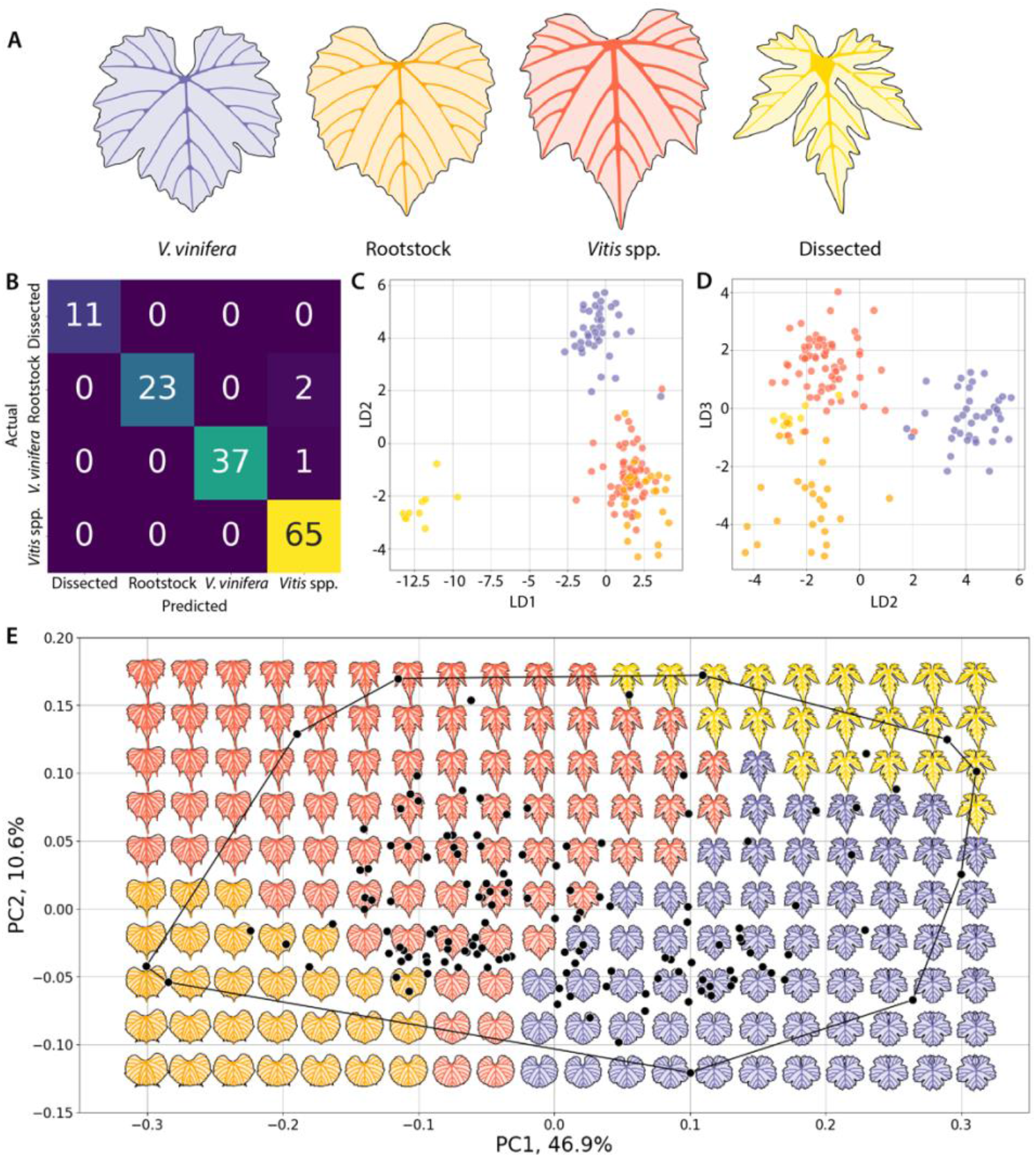
Linear Discriminant Analysis (LDA) model by genotype. **A)** Generalized Procrustes Analysis mean leaves by genotype levels, indicated by color and text: *V. vinifera*, Rootstock, *Vitis* spp., and Dissected. **B)** Confusion matrix plotting actual by predicted categories with numbers of each class and indicated by color. **C)** Data colored by genotype projected onto a plot of LD2 vs. LD1 and **D)** LD3 vs. LD2. **E)** Principal Component Analysis (PCA) morphospace. Eigenleaf representations are colored by genotype prediction using the LDA classifier. Data is plotted and bounded by a convex hull.

**Figure 6:**
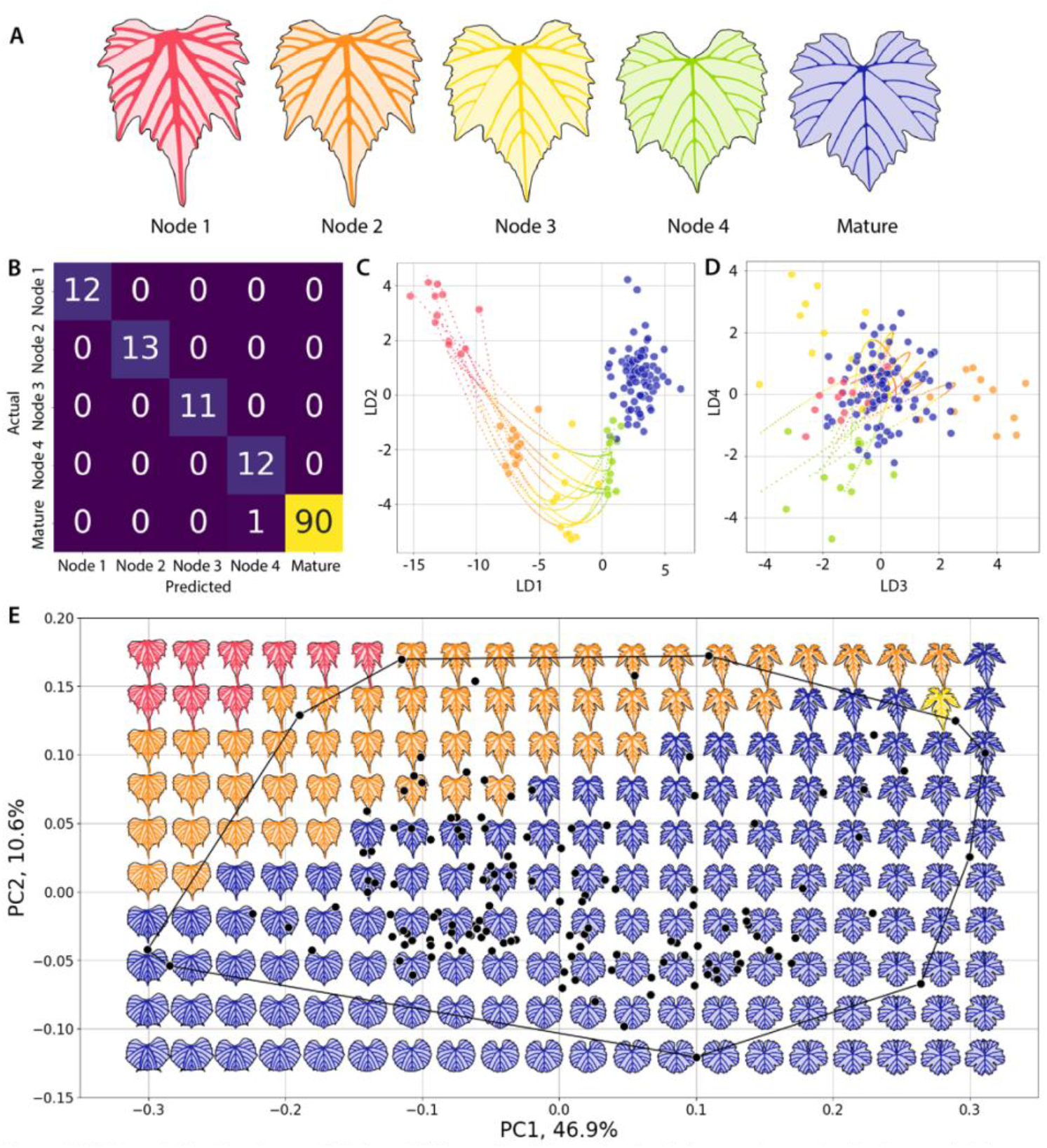
LDA model by developmental stage. **A)** Generalized Procrustes Analysis mean leaves by developmental stage, indicated by color and text: node 1, node 2, node 3, node 4, and mature. **B)** Confusion matrix plotting actual by predicted categories with numbers of each class and indicated by color. **C)** Data colored by developmental stage and projected onto a plot of LD2 vs. LD1 and **D)** LD4 vs. LD3. Continuous developmental models for 10 species from node 1 to node 4 and their predicted developmental stage are indicated by smaller points. **E)** PCA morphospace. Eigenleaf representations are colored by developmental stage prediction using the LDA classifier. Data is plotted and bounded by a convex hull.

## Discussion

Ampelography relies on trained, human cognition to identify corresponding features that vary by variety (Bodor-Pesti et al., 2023). Although human-derived ampelographic data is highly accurate, it is constrained by the time to manually measure it: either thousands of leaves using dozens of landmarks or hundreds of leaves with thousands of landmarks (as in this work) can be measured (**Table 1**).

Although we automate the detection of landmarks and veins from trace data, truly automatic detection of ampelographic features remains elusive. Not only does background noise prevent accurate classification of vein and blade pixels, but occlusion (for example, overlapping lobes) prevents detecting key features. We note that the vein and blade traces we produce on raw images are the perfect training set for automatic feature detection by machine learning models. Contrastingly, machine learning and convolutional neural networks can achieve high classification rates directly from images (Magalhães et al., 2023). They also overcome problems associated with noise and leverage high numbers of *de novo* features. However, while machine learning can achieve high classification rates and requires little data manipulation to use, it is difficult to interpret the features used for classification, much less within an established ampelographic framework that can be extrapolated to other contexts.

Previously, using thousands of leaves with only 21 landmarks (Chitwood and Mullins, 2022), we similarly sampled the *Vitis* morphospace as we have done here. Consistent between both works, genetic and developmental contributions to leaf shape are orthogonal and can be predicted independently of the other. Unlike our previous work, by sampling both sides of the leaf and with a saturating number of pseudo-landmarks, the resulting morphospace is defined by realistic eigenleaf representations. We can sample this space beyond just the first two PCs to create synthetic leaves that represent morphological diversity within *Vitis*. As an example, we consider the first 9 PCs representing 88% of all shape variation (**Fig. 4B**). To estimate the borders of this 9 dimensional space, we can calculate a convex hull and uniformly sample synthetic leaves within it (**Fig. 7**).

**Figure 7:**
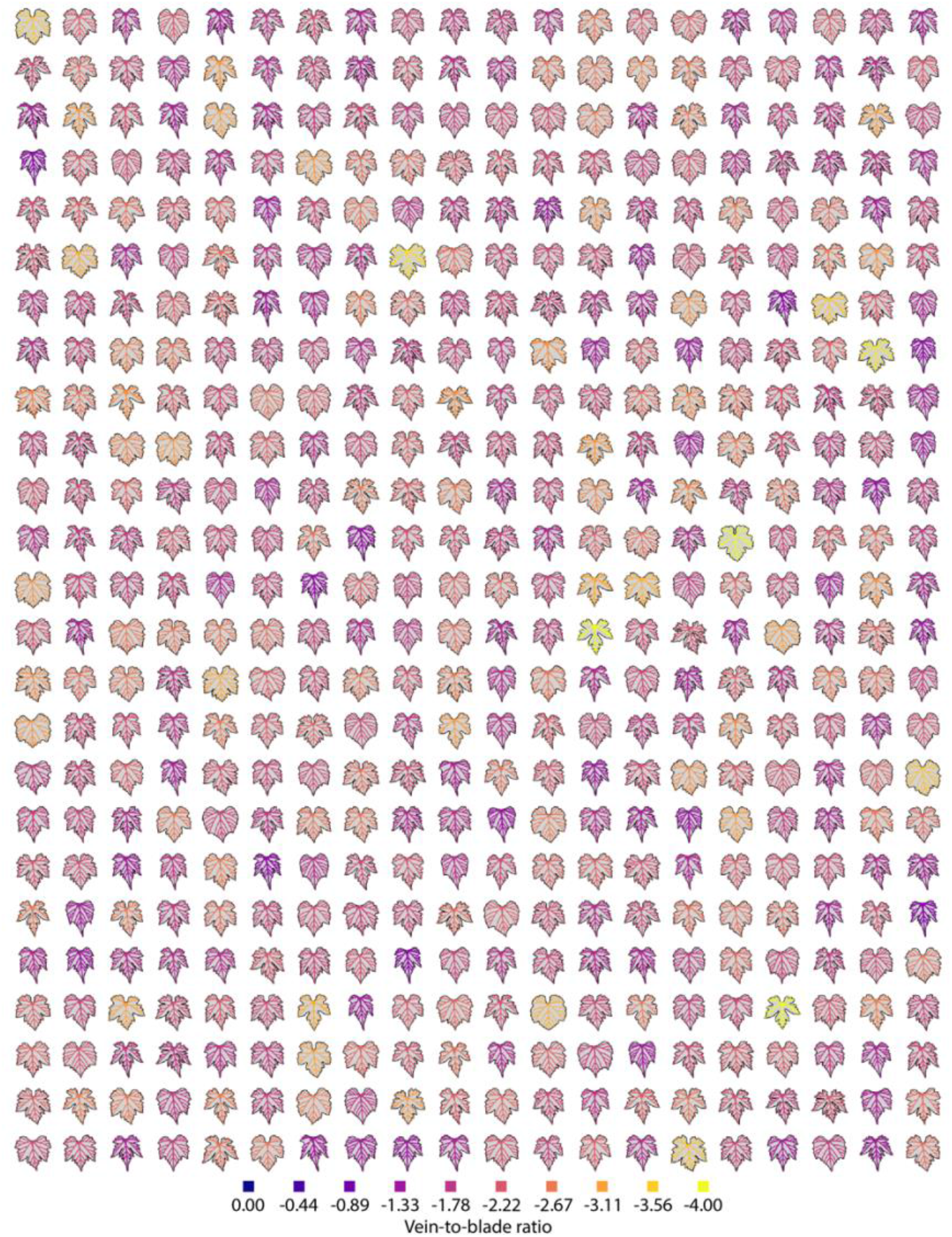
Synthetic leaves. Using the inverse transform of the morphospace on uniform sampling of a convex hull of the data calculated using the first nine principal components, 500 synthetic leaves are shown. Vasculature is colored by the natural log of the vein to blade area ratio as indicated.

Each of these synthetic leaves is unique and represents a position within the genetic and developmental boundaries of the *Vitis* morphospace. Each is a theoretical representation of a possible grapevine leaf, the features of which are already defined.

Such synthetic leaves offer new opportunities in data analysis. Because they are sampled from within the boundaries of a known morphospace, it is possible to predict and visualize leaves we have not sampled. For example, in our own data we have yet to sample developmental series from *V. vinifera* varieties the same way we have for *Vitis* spp. Yet, because we can estimate developmental variation in relation to variation that defines differences between species and varieties (**Figs. 4-6**), we look forward to predicting developmental series of wine and table grape varieties and comparing them to empirically sampled leaves, as we previously proposed (Chitwood and Mullins, 2022). Another possibility is to bypass manual measurements by projecting onto a precalculated morphospace. Because synthetic leaves already contain most the features found in grapevine leaves, subsets of features can be projected back onto the high resolution morphospace. For example, pseudo-landmarks around the blade outline using tip and petiolar junction landmarks, or the large number of leaves we have previously landmarked using 21 points, can be calculated in synthetic leaves as well. The best match between an empirically sampled leaf with a synthetic leaf can then be found using the Procrustes distance between the two and projected back into the high resolution space.

Finally, because synthetic leaves are realistic and capture intricate features of leaf morphology, they could be used as inputs into convolutional neural network classifiers to increase robustness and accuracy by augmenting and broadening training data.

## Conclusion

The ability to create realistic, synthetic leaves with intricate features arises from the constrained morphology of grapevine leaves with numerous homologous features that lends itself to morphometric analysis. While manual ampelographic measurements and convolutional neural network approaches are forward approaches that detect features first to classify leaves, the approach we describe here is reverse: by estimating an underlying morphospace that can predict synthetic leaves, we can use such theoretical leaves to anticipate the leaves we have yet to sample and develop tools that increase our abilities to classify varieties and learn about the underlying biology of grapevines.

## Acknowledgements

This work was supported by National Science Foundation Plant Genome Research Program award numbers IOS-2310355, IOS-2310356, and IOS-2310357.

## Author Contributions

Data collection: DHC, ETL, MGFA, SA, SWB, DPC, EEC, AJE, MFF, QF, ESH, CH, JJ, BMK, NSK, AL, BLM, JTP, WLGP, SIR, SER, FLS, YS, CCS, MKS, CSS, PW, JW, LDG Student advising and supervision: DHC, ETL, LDG Conceptualization: DHC, LDG Data analysis: DHC, ETL, LDG Writing: DHC Reading and revising: DHC, ETL, MGFA, SA, SWB, DPC, EEC, AJE, MFF, QF, ESH, CH, JJ, BMK, NSK, AL, BLM, JTP, WLGP, SIR, SER, FLS, YS, CCS, MKS, CSS, PW, JW, LDG

## Data Availability Statement

The data that support the findings of this study are openly available in github at https://github.com/DanChitwood/synthetic_leaves

## Conflict of Interest Statement

The authors report no conflict of interest.

## Notes

https://github.com/DanChitwood/synthetic_leaves

### Competing Interest Statement

The authors have declared no competing interest.

## References

Bodor-Pesti, P., Taranyi, D., Deák, T., Nyitrainé Sárdy, D.Á. & Varga, Z. (2023). A Review of Ampelometry: Morphometric Characterization of the Grape (Vitis spp.) Leaf. Plants, 12(3), 452.

Braden, B. (1986). The surveyor’s area formula. The College Mathematics Journal, 17(4), 326–337.

Bryson, A.E., Wilson Brown, M., Mullins, J., Dong, W., Bahmani, K., Bornowski, N., Chiu, C., Engelgau, P., Gettings, B., Gomezcano, F. Gregory, L.M., et al. (2020). Composite modeling of leaf shape along shoots discriminates Vitis species better than individual leaves. Applications in Plant Sciences, 8(12), e11404.

Chitwood, D.H., Ranjan, A., Martinez, C.C., Headland, L.R., Thiem, T., Kumar, R., Covington, M.F., Hatcher, T., Naylor, D.T., Zimmerman, S. Downs, N., et al. (2014). A modern ampelography: a genetic basis for leaf shape and venation patterning in grape. Plant physiology, 164(1), 259–272.

Chitwood, D.H. & Sinha, N.R. (2016). Evolutionary and environmental forces sculpting leaf development. Current Biology, 26(7), R297–R306.

Chitwood, D.H., Klein, L.L., O’Hanlon, R., Chacko, S., Greg, M., Kitchen, C., Miller, A.J. & Londo, J.P. (2016a). Latent developmental and evolutionary shapes embedded within the grapevine leaf. New Phytologist, 210(1), 343–355.

Chitwood, D.H., Rundell, S.M., Li, D.Y., Woodford, Q.L., Yu, T.T., Lopez, J.R., Greenblatt, D., Kang, J. & Londo, J.P. (2016b). Climate and developmental plasticity: interannual variability in grapevine leaf morphology. Plant physiology, 170(3), 1480–1491.

Chitwood, D.H. (2020). Data from: Vitis vinifera leaf images [Dataset]. Dryad. 10.5061/dryad.g79cnp5mn

Chitwood, D.H., VanBuren, R., Migicovsky, Z., Frank, M., Londo, J. (2020). Data from: Latent developmental and evolutionary shapes embedded within the grapevine leaf [Dataset]. Dryad. 10.5061/dryad.zkh189377

Chitwood, D.H. (2021). The shapes of wine and table grape leaves: An ampelometric study inspired by the methods of Pierre Galet. Plants, People, Planet, 3(2), 155–170.

Chitwood, D.H., Mullins, J., Migicovsky, Z., Frank, M., VanBuren, R. & Londo, J.P. (2021). Vein‐to‐blade ratio is an allometric indicator of leaf size and plasticity. American Journal of Botany, 108(4), 571–579.

Chitwood, D.H. and Mullins, J. (2022). A predicted developmental and evolutionary morphospace for grapevine leaves. Quantitative Plant Biology, 3, e22.

Chitwood, D.H. (2024). Data from: A high resolution model of the grapevine leaf morphospace predicts synthetic leaves [Dataset]. figshare. Figure. 10.6084/m9.figshare.25325980.v1

Galet, P. (1979). A Practical Ampelography (L.T. Morton, Trans.). Ithaca, USA: Cornell University 522 Press.

Galet, P. (1985). Précis d’ampélographie pratique, 5 ed., Montpellier, France: Déhan.

Galet, P. (1988). Cépages et vignobles de France, vol. I, Les vignes américaines. Montpellier, France: Déhan.

Galet, P. (1990). Cépages et vignobles de France, vol. II. L’ampélographie française. Montpellier, France: Déhan.

Galet, P. (2000). Dictionnaire encyclopédique des cépages. Paris, France: Hachette

Goethe, H. (1876). Note sur l’ampelographie. Congress of Marburg, September 18.

Goethe, H. (1878). Handbuch der Ampelographie. Austria, Graz: Commission-Verlag von Leykam Josefsthal.

Goodall, C. (1991). Procrustes methods in the statistical analysis of shape. Journal of the Royal Statistical Society: Series B (Methodological), 53(2), 285–321.

Gower, J.C. (1975). Generalized procrustes analysis. Psychometrika, 40, 33–51.

Harris, C.R., Millman, K.J., Van Der Walt, S.J., Gommers, R., Virtanen, P., Cournapeau, D., Wieser, E., Taylor, J., Berg, S., Smith, N.J. & Kern, R. (2020). Array programming with NumPy. Nature, 585(7825), 357–362.

Hunter, J.D. (2007). Matplotlib: A 2D graphics environment. Computing in Science & Engineering, 9(03), 90–95.

Kluyver, T. et al./person-group>. (2016). Jupyter Notebooks – a publishing format for reproducible computational workflows. In F. Loizides & B. Schmidt, eds. Positioning and Power in Academic Publishing: Players, Agents and Agendas. 87–90.

Magalhães, S.C., Castro, L., Rodrigues, L., Padilha, T.C., De Carvalho, F., Dos Santos, F.N., Pinho, T., Moreira, G., Cunha, J., Cunha, M. & Silva, P. (2023). Toward grapevine digital ampelometry through vision deep learning models. IEEE Sensors Journal, 99(1), 1.

Martínez MC & Grenan S. (1999). A graphic reconstruction method of an average vine leaf. Agronomie, EDP Sciences 19(6): 491–507.

McKinney, W. & Team, P.D. (2015). Pandas-Powerful python data analysis toolkit. Release 0.22.0.

Meister, A.L.F. 1769. Generalia de genesi figurarum planarum et inde pendentibus earum affectionibus.

Pedregosa, F., Varoquaux, G., Gramfort, A., Michel, V., Thirion, B., Grisel, O., Blondel, M., Prettenhofer, P., Weiss, R., Dubourg, V. & Vanderplas, J. (2011). Scikit-learn: Machine learning in Python. The Journal of Machine Learning Research, 12, 2825–2830.

Ravaz, L. (1902). Les vignes américaines: Porte-greffes et producteurs directs. Goulet, Montpellier and Paris. Digitized by Google Books from Cornell University.

Schindelin, J., Arganda-Carreras, I., Frise, E., Kaynig, V., Longair, M., Pietzsch, T., Preibisch, S., Rueden, C., Saalfeld, S., Schmid, B. & Tinevez, J.Y. (2012). Fiji: an open-source platform for biological-image analysis. Nature Methods, 9(7), 676–682.

Schneider, C.A., Rasband, W.S. & Eliceiri, K.W. (2012). NIH Image to ImageJ: 25 years of image analysis. Nature Methods, 9(7), 671–675.

Virtanen, P., Gommers, R., Oliphant, T.E., Haberland, M., Reddy, T., Cournapeau, D., Burovski, E., Peterson, P., Weckesser, W., Bright, J. & Van Der Walt, S.J. (2020). SciPy 1.0: fundamental algorithms for scientific computing in Python. Nature Methods, 17(3), 261–272.

Waskom, M.L., 2021. Seaborn: statistical data visualization. Journal of Open Source Software, 6(60), 3021.

